# StoichRNA: A Scalable Framework to Map and Optimize Combinatorial RNA Systems at Single-Cell Resolution

**DOI:** 10.64898/2026.10.19.760129

**Authors:** Qiaochu Liu, Lei Wang

## Abstract

RNA-based genetic cargos have emerged as versatile tools in both fundamental research and translational medicine, with growing applications in regenerative medicine, gene therapy, cell therapy, and vaccination. As the field advances beyond single-RNA delivery, coordinated engineering of multiple RNA species is enabling increasingly sophisticated and combinatorial biological functions. However, this shift raises a fundamental challenge: what combination and stoichiometry of RNA species should be delivered together to achieve a desired cellular function? As the number of RNA components increases, the design space expands combinatorially, making conventional optimization strategies labor-intensive and poorly scalable. Here, we introduce StoichRNA, a framework for scalable exploration of multi-RNA stoichiometry in a single experiment. Individual RNA species are delivered independently, each coupled to a distinct fluorescent marker that reports its intracellular abundance. Because uptake is stochastic, each cell acquires a unique combination and copy number of RNA components. Across a sufficiently large population, cells collectively sample a broad stoichiometric landscape, allowing single-cell readouts such as flow cytometry to quantitatively map system output as a function of intracellular RNA composition. We first establish the conceptual validity of this approach by showing that StoichRNA-derived single-cell data can recover and predict the behavior of fixed-ratio RNA formulations. We then apply this framework to representative RNA circuits, demonstrating that StoichRNA can reveal design principles and identify improved stoichiometries that would be difficult to uncover through conventional workflows. Finally, we extend StoichRNA optimization to translationally relevant settings, including control of cell-state engineering in human induced pluripotent stem cells and tuning of RNA-based logic circuits for precision medicine applications. Together, these results establish StoichRNA as a generalizable strategy for engineering combinatorial RNA systems and provide a scalable solution to a central bottleneck in RNA circuit design.

## INTRODUCTION

RNA-based genetic cargos have emerged as a powerful and versatile class of tools in both fundamental research(1) and translational medicine(2), with growing applications in regenerative medicine(3), gene therapy(4), cell therapy(5), and vaccination(6). As these technologies continue to mature, the field is increasingly moving beyond single-RNA delivery toward coordinated action of multiple RNA species engineering to enable more sophisticated and combinatorial biological functions(7). Many emerging RNA applications inherently depend on the concerted action of multiple RNA components. For example, efficient genome editing requires delivery of mRNAs encoding CRISPR-associated proteins together with their corresponding guide RNAs (8). Similarly, synthetic mammalian RNA circuits increasingly rely on multiple interacting transcripts, making circuit performance sensitive to the relative abundance of each component (9–11). In vaccine development, co-delivery of mRNAs encoding multiple viral antigens has been shown to elicit broader and more protective immune responses than single-antigen approaches(12). Together, these examples highlight a broader transition in RNA technology: from single-agent delivery toward the engineering of combinatorial RNA systems.

A central challenge in combinatorial RNA systems is that functional output depends not only on the identity of individual components, but also on their relative intracellular stoichiometry. Conventional co-delivery of separately formulated RNAs can produce substantial cell-to-cell variability in component ratios, motivating increasing interest in co-encapsulation strategies that deliver multiple RNA species at defined proportions within the same carrier (13). However, this creates a fundamental design challenge: which RNA components, and in what relative ratios, should be combined to achieve optimal function? As the number of components increases, the space of possible combinations and stoichiometries expands rapidly, making conventional condition-by-condition optimization labor-intensive and poorly scalable.

Addressing this challenge requires systematic exploration of multicomponent stoichiometry rather than characterization of individual parts in isolation. Recent studies of mammalian genetic circuits have begun to tackle this problem using approaches such as poly-transfection(14), programmable multigene transcription systems(15), resource-aware circuit design(16), and large-scale combinatorial screening(17). Collectively, these studies demonstrate that relative component abundance and competition for cellular resources can substantially alter circuit performance. However, these approaches have primarily been developed for DNA-encoded circuits, in which component levels are specified through promoter strength, construct architecture, or DNA copy number. Among them, the poly-transfection strategy is particularly attractive as a foundation for adaptation to RNA systems because it enables systematic exploration of multicomponent stoichiometry using transient transfection and flow cytometry based readouts, without requiring stable genomic integration or next-generation sequencing(14). This experimentally accessible framework can therefore be readily implemented in a standard laboratory setting and provides a natural starting point for direct optimization of RNA mixtures. Nevertheless, direct translation to RNA requires additional considerations because RNA is governed by distinct delivery properties(18), regulatory mechanisms (19), chemical modifications (20), and expression kinetics (21). In vitro-transcribed RNA lacks promoter-based transcriptional control exhibits rapid and transient expression and can be further shaped by RNA-specific sequence and chemical features (21). These differences necessitate a dedicated framework for directly optimizing multi-RNA stoichiometry at the RNA level.

Here, we introduce a StoichRNA transfection framework that enables scalable exploration of multi-RNA stoichiometry in a single experiment. In this approach, individual RNA species are delivered independently, each coupled with a distinct fluorescent marker that reports its intracellular abundance (quantitatively traceable). Because uptake is stochastic, each cell acquires a unique combination and copy number of RNA components. Across a sufficiently large population, cells in one transfection collectively sample a broad distribution of stoichiometric states, such that each cell effectively serves as an independent reaction unit within a high-dimensional combinatorial design space. By coupling this sampling strategy with single-cell readouts such as flow cytometry, StoichRNA quantitatively maps system output as a function of intracellular RNA composition. In this way, StoichRNA converts stochastic delivery from a source of variability into a scalable search mechanism for identifying optimal RNA combinations and ratios without exhaustive condition-by-condition testing. We first established the conceptual validity of this approach by benchmarked StoichRNA against multiple fixed-ratio co-transfections, showing that StoichRNA-derived single-cell data can recover and predict the behavior of fixed-ratio RNA formulations. We next tested the hypothesis that DNA- and RNA-encoded formats of the same circuit do not share an optimal stoichiometry. We then apply this framework to representative RNA circuits, demonstrating that StoichRNA can reveal design principles and identify improved stoichiometries that would be difficult to uncover through conventional workflows. Finally, we extend StoichRNA optimization to translationally relevant settings, including control of cell-state engineering in human induced pluripotent stem cells and tuning of RNA-based logic circuits for precision medicine applications. Together, these results establish StoichRNA as a generalizable strategy for engineering combinatorial RNA systems and provide a scalable solution to a central bottleneck in RNA circuit design.

## MATERIALS AND METHODS

### DNA assembly framework

The plasmids used in this study are constructed under a standardized Golden Gate assembly method which has been published previously (14, 22). First, Golden Gate assembly was used to construct plasmids for each transcription unit (termed plasmid level 1 or pL1 plasmids) from a number of basic input parts (termed plasmid level 0 or pL0) including insulators (pL0-I), promoters (pL0-P), 5’ untranslated regions (UTRs) (pL0-5), gene coding sequences (pL0-G), 3’ UTRs (pL0-3), and transcriptional terminators/poly-A sequences (pL0-T). The forming transcription units can be then cloned into corresponding position backbones (pL1-BB) with ST1-2, ST2-3 and ST3-4X for positions 1, 2 and 3, allowing multiple transcription units to be assembled into one plasmid for further experiment.

### RNA *In vitro* transcription

The plasmid used to generate mRNA is assembled with a T7 promoter with a modified transcription initiation sequence of AG at the 5’ UTR (pL0-5) location and a 140bp long poly-A tail at the terminator location (pL0-T). A BsmBI enzyme cutting site is designed at the end of the poly-A tail for DNA plasmid linearization. The *in vitro* transcription is carried out using HiScribe® T7 mRNA Kit with CleanCap® Reagent AG (New England Biolabs), with Pseudo-UTP (New England Biolabs) replacing the UTP in the kit. Then the synthesized mRNA transcript is purified using RNA purification kit (Zymo research), following the manufacturer’s protocol.

### Cell culture

The HEK-293T cell line used in this study was obtained from Thermo Fisher. This cell line is maintained in DMEM containing 4.5 g/l glucose, L-glutamine, and sodium pyruvate (Corning) which was supplemented with 10% FBS (Corning) and were grown at 37°C and 5% CO2.

The PGP1 hiPS cells were a kind gift from George Church (Harvard University) and can be obtained from Coriell. The hiPS cells were cultivated under sterile conditions in mTeSR Plus (StemCell Technologies, Vancouver). Tissue culture plates were coated for 1 h at room temperature with Matrigel (Corning) diluted according to the manufacturer’s instructions in ice-cold DMEM/F-12 with 15 mM HEPES medium (StemCell Technologies, Vancouver). Routine passaging was performed by incubating hiPS cells for 7 min in Gentle Cell Dissociation Reagent (StemCell Technologies, Vancouver) at 37 °C followed by three 2 ml washes in DMEM/F-12 medium and mechanical dissociation (no. 3010 cell scraper, Corning). Subsequently, the cells were taken up in 5 ml of DMEM/F-12, centrifuged at 500 r.p.m. for 3 min and resuspended in mTeSR Plus. The clump size was assessed by eye and if necessary, a further reduction in size was performed by gently pipetting the suspension. Single-cell suspensions were generated by incubating hiPS cell colonies for 5 min in Accutase (StemCell Technologies, Vancouver) at 37 °C, subsequently resuspending this single-cell solution in 5 ml of DMEM/F-12 medium containing 10 μM Y-27632 dihydrochloride (Tocris Biosciences) solubilized in cell culture grade dimethylsulfoxide (Sigma-Aldrich), centrifuging it at 500 r.p.m. for 3 min and resuspending the pellet in mTeSR Plus or DMEM/F-12 supplemented with Y-27632 at a final concentration of 10 μM for counting. All cell lines were grown at 37 °C and 5% CO2. All cell lines were tested negative for mycoplasma.

### Transfections

All HEK293T cell transfections were performed using Lipofectamine MessengerMax reagent (Thermal Fisher) following manufacturer’s protocol. For hiPSC, transfections were performed using Lipofectamine Stem regent (Thermal Fisher) following manufacturer’s protocol. All co-transfections are done in 24 well plate format with 500ng total mRNA.100,000 HEK293T or 50,000 hiPSC cells are transfected per well. For StoichRNA experiments, each mRNA transfection complex was prepared separately. In brief, to set up a two complexes StoichRNA reaction in one well of the 24 well plate; mRNA 1 is mixed with 0.5 uL of transfection reagent; mRNA 2 is mixed with 0.5 uL of transfection reagent separately. Each transfection complexes are allowed to form following the manufacture’s protocol. The 2 complexes are then added as liquid droplet at different regions of the well without touching each other. Cells are then seeded on top of the 2 separate transfection complexes. For each transfection complexes; 200-300ng of mRNA are mixed with 0.5uL of MessengerMAX reagents and diluted into a total of 25uL with Opti-MEM.

### Flow cytometry

Flow cytometry analysis data in this study were collected through a Beckman Coulter CytoFLEX S cytometer located in the Institute of Chemical Imaging of Living Systems at Northeastern University. The cytometer is equipped with a 405 nm laser with 450/45 nm filter (‘PB450’) for measuring BFP, 488 nm laser with 525/40 filter (‘FITC’) for measuring GFP, 561 nm laser with 610/20 nm filter (‘ECD’) for measuring RFP and 640 nm laser with 712/25 nm filter (‘APC’) for measuring iRFP (iRFP720). For each co-transfection, at least 20,000 cells/events were collected per sample. For StoichRNA experiment with 2 complexes, 100,000 cells are/events collected, and 400,000 cells/events are collected for 3 complexes experiment. All data files were saved, and the gating and compensation were applied after data collection.

### Data analysis

We have developed a Python pipeline to facilitate analysis of multi-dimensional StoichRNA experiment data. Code may be found on Github.

### Statistical analysis

Statistical analysis of this work is based on the log-normal distribution of the fluorescence mRNA expressions within the cells.

## RESULTS

### StoichRNA enables one-pot exploration of multi-RNA stoichiometry

To enable parallel exploration of RNA component ratios, we developed StoichRNA, an RNA-level implementation of the poly-transfection principle previously established for plasmid DNA(14). In conventional RNA co-transfection, different RNA components are mixed at a defined input ratio before transfection complexes are formed, favoring coupled delivery and correlated expression across cells (Fig. 1A). StoichRNA instead uses separately prepared transfection complexes for each RNA component (Fig. 1B). These complexes were deposited in different regions of the same cell culture well, followed by cell addition. This design exploits variation in the uptake of the separately prepared complexes to generate diverse combinations of RNA component levels within a single cell population by a single transfection.

**Figure 1.**
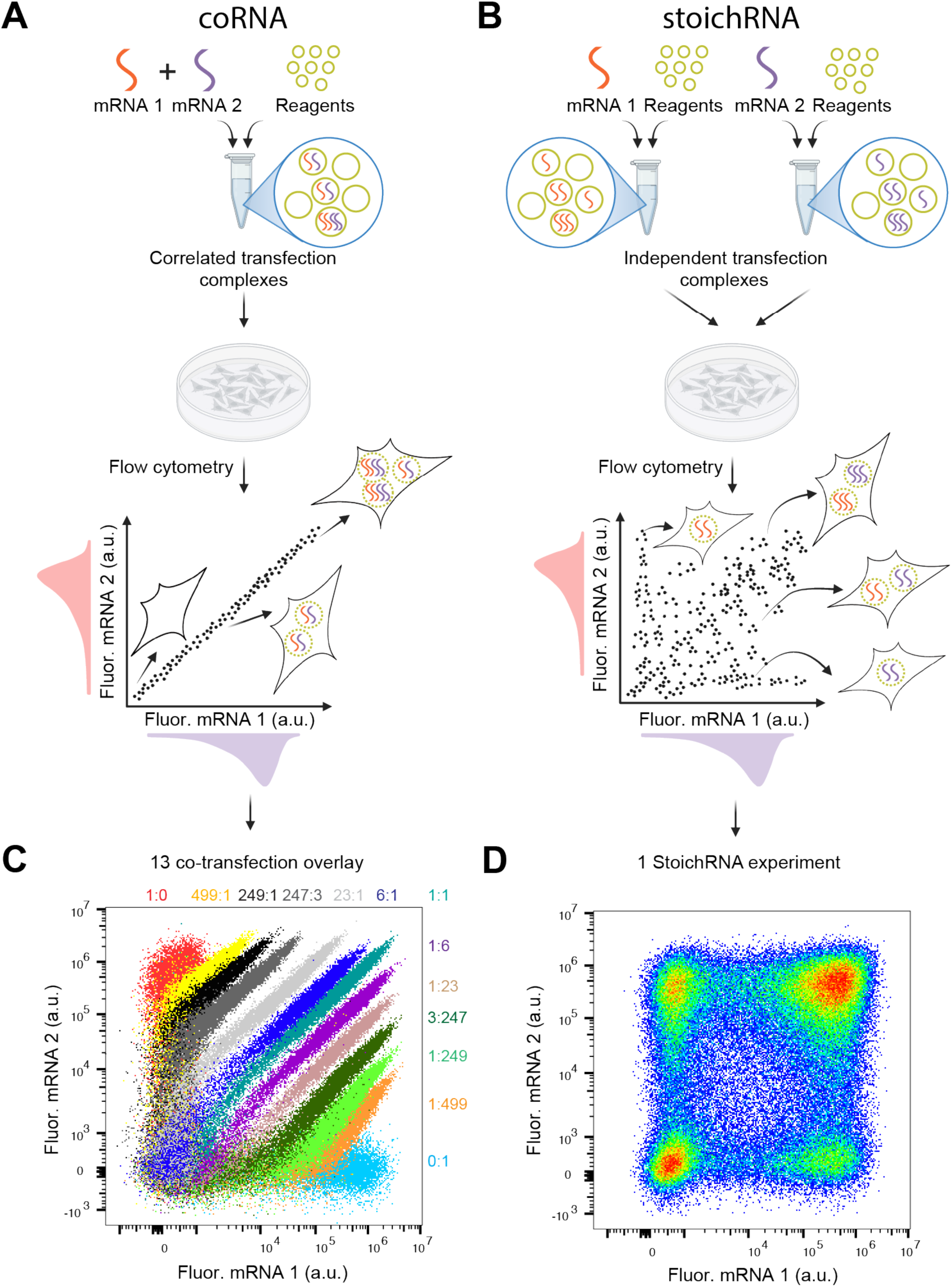
Overview of conventional coRNA and StoichRNA transfections. (A) coRNA: Two fluorescent mRNAs are mixed before combined with transfection reagent. Resulting in transfection complex encapsulates zero, few, or many copies of both mRNA 1 and mRNA 2, with the ratio of the two mRNA species within a complex is set by the input mix and is therefore approximately constant across complexes. A given cell may take up zero, few, or many complexes, yet because every complex carries the same ratio of two mRNA species, the relative fraction of mRNA 1 to mRNA 2 inside each cell stays effectively unchanged. The result is strongly correlated expression of the two fluorescent mRNAs by flow cytometry, with cells distributed along a single diagonal line. (B) StoichRNA: Each fluorescent mRNA component is mixed with transfection reagent separately, so every complex contains only one mRNA species. The single-species complexes are then delivered to cells simultaneously. An individual cell independently samples complex 1 and complex 2, so the uptake of the two species is stochastic and uncorrelated, generating a population in which the mRNA 1: mRNA 2 varies continuously across population and spans a wide range. (C): Flow cytometry scatter plot of 13 different ratios of mRNA1: mRNA 2 overlay. (D): Flow cytometry scatter plot of 1 StoichRNA experiment of mRNA1 and mRNA 2.

We first evaluated this sampling strategy using two mRNAs encoding distinct fluorescent proteins and measured their expression levels by flow cytometry. Conventional co-transfection produced ratio-dependent single-cell distributions: conditions containing both mRNAs generally formed diagonal bands, reflecting correlated expression of the two reporters. Changing the input RNA ratio shifted the distribution, such that sampling different relative expression levels required a series of independently prepared formulations (Fig. 1C). In contrast, a single StoichRNA transfection generated a broad two-dimensional fluorescence distribution, including cells with high expression of either reporter relative to the other and cells co-expressing both reporters at different levels (Fig. 1D). Thus, StoichRNA sampled a wide range of relative reporter-expression levels within one transfection rather than concentrating cells along the correlated distributions associated with individual co-transfection formulations.

This broad sampling provides the experimental basis for characterizing RNA circuits across multiple component combinations without separately preparing each input ratio. For subsequent circuit experiments, each functional RNA component was paired with a distinct fluorescent-marker RNA during complex formation. Joint measurement of marker fluorescence and circuit output allowed circuit behavior to be analyzed across marker-defined component levels, providing a basis for identifying optimal-performing regions of RNA stoichiometric space.

### Nucleic acid format and RNA chemistry reshape circuit performance landscapes

We next asked whether the optimal performance and corresponding stoichiometric condition of a circuit are preserved between plasmid DNA and in vitro-transcribed RNA implementations. To test this, we used a CasE based OFF circuit comprising two modules: CasE expression module, and a GFP reporter module carrying a CasE target site in the 5′ UTR. CasE cleaves the reporter transcript at this site, repressing GFP expression.

For the DNA implementation, each circuit module was paired with a constitutively expressed fluorescent transfection marker, with BFP tracking the CasE module and iRFP tracking the reporter module (Fig. 2A). Cells were binned by the two marker signals to map fold change across the sampled expression space. DNA poly-transfection yielded a maximum fold change of approximately 56.4x (Fig. 2D), with the optimal performing region occurring at high reporter (iRFP) and intermediate CasE (BFP) plasmid levels.

**Figure 2.**
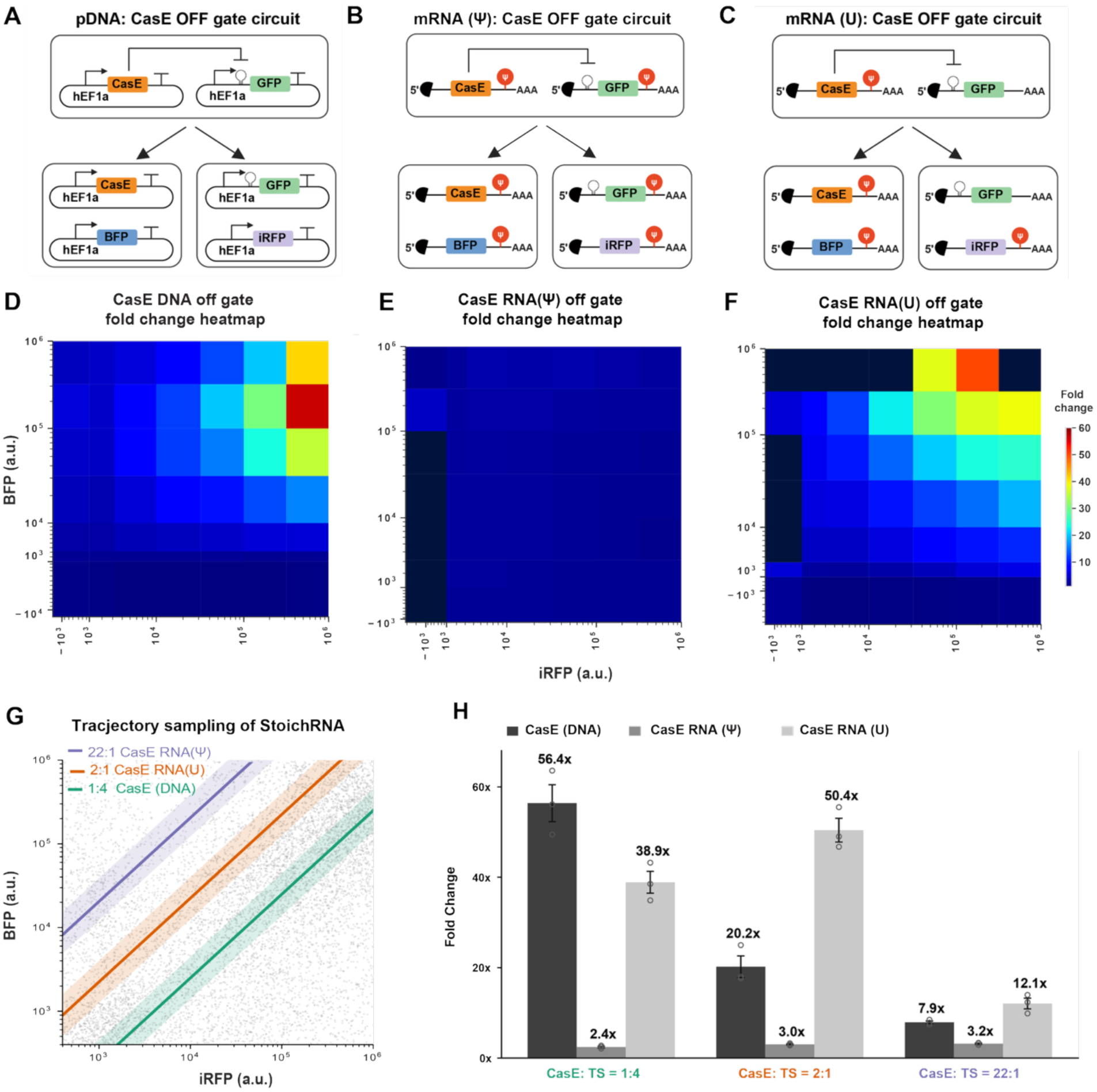
StoichRNA reveals nucleic acid format and RNA chemistry reshape circuit performance landscapes. (A) Circuit schematic of the plasmid DNA-encoded CasE OFF gate. BFP plasmid is paired with the constitutive CasE expression plasmid to report CasE expression, and iRFP plasmid is paired with the CasE circuit reporter (GFP output). (B) Circuit schematic of the mRNA-encoded CasE OFF gate. Marker and circuit component pairing is identical to A, implemented in mRNA format. All circuit components are Ψ-substituted. (C) Circuit schematic of the mRNA-encoded CasE OFF gate as in B, except that the CasE reporter mRNA is unmodified (uridine, not Ψ-substituted). (D) Fold change heatmap for the plasmid DNA CasE system. Cells were binned into an [7 × 7] grid of biexponentially spaced bins across iRFP (x) and BFP (y) marker fluorescence, each axis spanning 10⁻³ to 10⁶ (a.u.); binning and axes are identical in D–F. Fold change is calculated per bin as the ratio of mean of output GFP fluorescence in the [without CasE control] to that in the [With CasE] condition, so that values >1 indicate repression. Maximum fold change (∼56.4×) occurs in the intermediate-BFP, high-iRFP region of marker space. (E) Fold change heatmap for the fully Ψ-substituted CasE mRNA system. Ψ-substituted CasE mRNA system is analyzed under the same bin analysis in D. Fold change remains between 2.0× to 3.2× across the entire marker space. (F) Fold change heatmap for the CasE mRNA system carrying an unmodified (uridine) reporter. System is analyzed under the same bin analysis in D Maximum fold change (∼50.4×) occurs in the high-BFP, high-iRFP region. (G) Optimal CasE: CasE-target-site ratios identified for the three systems by trajectory sampling: 1:4 for the plasmid DNA system (green), 2:1 for CasE(U) (orange), and 22:1 for CasE(Ψ)(purple). (H) Fold-change histograms for the three systems at their respective optimal ratios; colors as in G.

We then implemented the same circuit using in vitro-transcribed RNA and profiled its performance using StoichRNA(Fig. 2B). We used pseudouridine (Ψ) substitution to enhance mRNA stability and protein expression while limiting innate immune activation (23–25). With a pseudouridine (Ψ)-substituted for both CasE encoding and reporter transcripts, the circuit exhibited severely attenuated repression across the whole sample space, with fold changes generally around 2.0x–3.2x (Fig. 2E). This predominantly low-fold-change landscape contrasted with the broad optimal performing region observed in the DNA implementation.

We hypothesized that Ψ substitution reduces the susceptibility of the reporter transcript to CasE cleavage, thereby weakening repression. To examine the contribution of reporter RNA modification, we next synthesized the reporter transcript with unmodified uridine while retaining a modified CasE-encoding transcript (Fig. 2C). This change restored strong repression, with fold changes of 50.4x being identified in the high-BFP/high-iRFP region (Fig. 2F). Thus, reporter RNA chemistry altered the circuit performance landscape without changing the circuit architecture. The recovery of repression with an unmodified reporter is consistent with the hypothesis that Ψ substitution reduces the susceptibility of the reporter transcript to CasE cleavage.

In addition to the heatmap analysis, we evaluated the optimal-performing conditions of the DNA and RNA implementations by computationally sampling trajectories across the sample space. Trajectory sampling allows StoichRNA data to be analyzed in a manner similar to conventional co-transfection: only cells falling close to a given trajectory are included, mimicking co-transfection at a fixed input ratio. The optimal CasE to target-site ratio was around 1:4 for the DNA implementation, 2:1 for the uridine RNA system, and 22:1 for the Ψ-modified RNA system, giving maximum fold changes of 56.4×, 50.4×, and 3.2×, respectively (Fig. 2G and H).

Together, these results show that CasE OFF-circuit performance depends on both marker-defined component levels and reporter RNA chemistry. They support direct characterization of RNA circuits rather than assuming that the operating conditions established in plasmid DNA circuits will translate to RNA circuits. StoichRNA enables these component-dependent responses to be mapped in one transfection for each RNA modification condition.

### Benchmarking StoichRNA vs fixed-ratio co-RNA transfection on RNA circuits

We next evaluated the expression space coverage and circuit output agreement of a single StoichRNA experiment relative to 13 fixed ratio co-transfections. We tested two widely used RNA-based OFF-gate circuits: one mediated by the RNA-binding protein L7Ae, and one by the endoribonuclease Csy4. In both systems, a repressor encoding mRNA was paired with a reporter mRNA carrying the cognate repressor target site in the 5’ UTR upstream of the GFP fluorescent output (Figure 3A, 3F). All repressor and reporter mRNAs were fully Ψ-substituted for both circuits. Repressor mRNA was tracked by a co-delivered BFP mRNA marker, and reporter mRNA by an iRFP marker.

**Figure 3.**
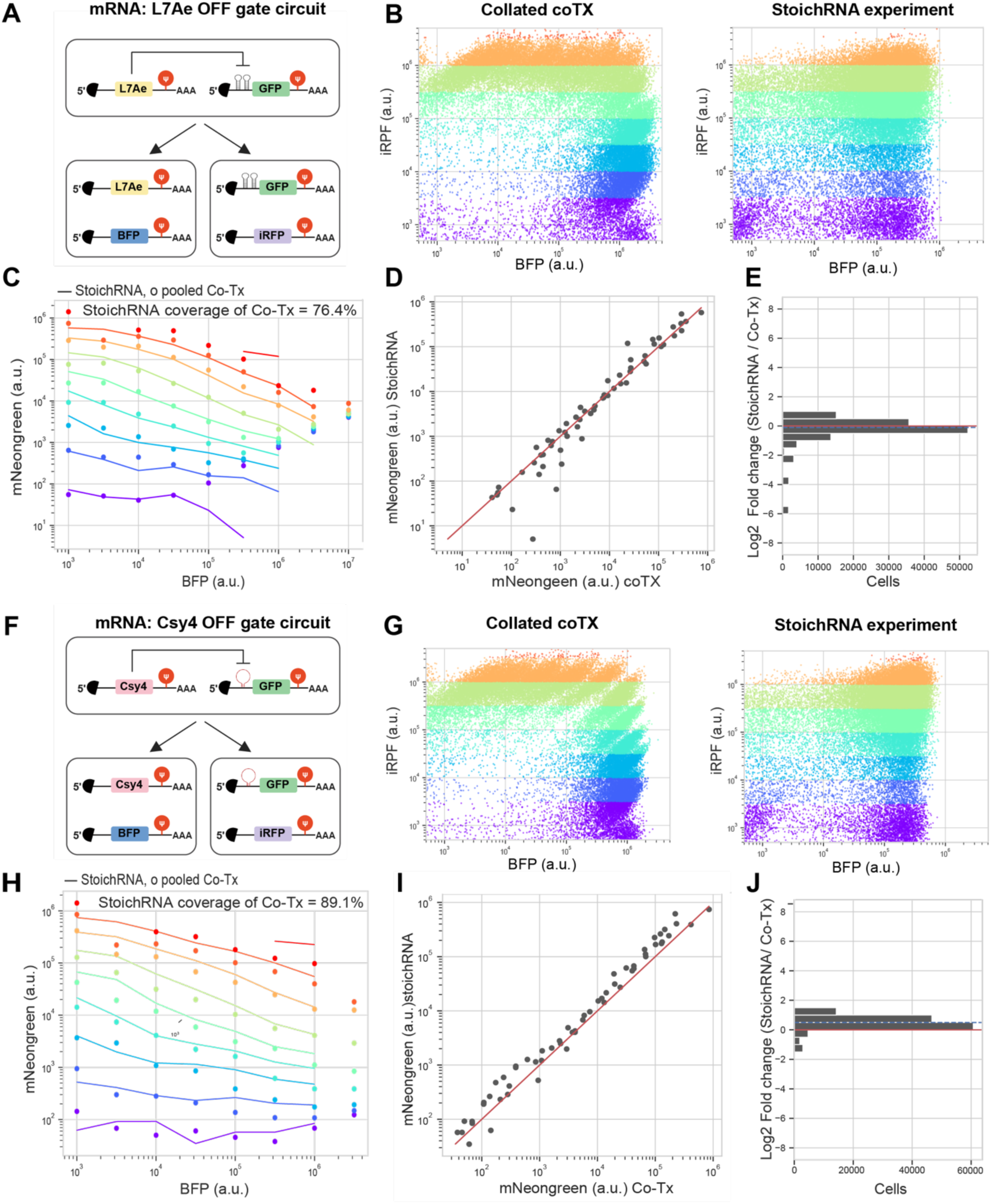
Benchmark RNA co-transfections with StoichRNA experiment. (A) Circuit schematic of the L7Ae system used for StoichRNA. BFP mRNA is paired with L7Ae mRNA to report L7Ae expression, and iRFP mRNA is paired with the circuit reporter (GFP output). All circuit components are Ψ-substituted. (B) 8 biexponentially spaced bins applied to the L7Ae system to stratify marker expression in the StoichRNA experiment. 13 co-transfections spanning different component ratios were pooled and binned identically for comparison with the matched StoichRNA experiment. Binning is illustrated using the BFP marker; bin colors are consistent in B, C, G and H. (C) Output coverage fit for the L7Ae system, comparing StoichRNA with collated co-transfections; y-axis, circuit output. Lines, StoichRNA; open circles, collated co-transfections. Colors denote the bins defined in B. (D) Correlation between StoichRNA output and the collated 13 RNA co-transfections for the L7Ae system. Each black point represents one bin; points fall near the red 1:1 line, indicating equivalent output between the two approaches. (E) Histogram of log2 fold change in output fluorescence between StoichRNA and collated co-transfections for the L7Ae system (mean = −0.175, SD = 0.800). A mean near zero and a narrow distribution indicate consistent readout between the two approaches. (F) Circuit schematic of the Csy4 system. The fluorescent marker configuration matches the L7Ae system in A. All circuit components are Ψ-substituted. (G) Bin analysis for the Csy4 system, as in B. (H) Output coverage fit for the Csy4 system, as in C. (I) Correlation between StoichRNA output and the collated 13 RNA co-transfections for the Csy4 system, as in D. (J) Histogram of log2 fold change in output fluorescence for the Csy4 system (mean = −0.441, SD = 0.461).

A single StoichRNA experiment sampled a broad region of the two-dimensional marker expression space that overlapped with the pooled co-transfected distributions. Data from these 13 co-transfections were collated into a single dataset for direct comparison with the corresponding StoichRNA experiment, and both datasets were binned by the two transfection-marker expression levels (Figure 3B, 3G). StoichRNA recovered 76.4% of L7Ae and 89.0% of Csy4 co-transfection bins (Figure 3C, 3H). Within the shared bins, circuit outputs from the StoichRNA broadly agreed with the collated co-transfections (Figure 3D, 3I), with log2 (StoichRNA/co-transfection) values of −0.179 for L7Ae and 0.441 for Csy4 (Figure 3E, 3J). Together, these results indicate that output values are broadly consistent either measured by StoichRNA or by a panel of co-transfections within the shared marker-expression space, supporting the utilization of StoichRNA for parallel characterization of RNA circuit responses.

### Optimizing a microRNA responsive RNA sensor using StoichRNA

Having benchmarked StoichRNA against fixed ratio RNA co-transfection, we next used StoichRNA to optimize a microRNA responsive RNA sensor. A microRNA-low sensor consists of a single mRNA encoding a fluorescent protein, flanked by two cassettes of four repeats of microRNA target sites with each located in the 5′ UTR and 3′ UTR. Upon delivery of the sensor RNA, the targeted endogenous microRNA binds the target sites, repressing reporter expression at high microRNA levels.

To demonstrate a microRNA-low sensor, we developed a hsa-miR-122-low sensor circuit in HEK293T cells, which do not express miR-122 endogenously (26), and used synthetic miR-122 mimics to span a range of input levels. Consistent with the previously tested transfection-marker layout, the miR-122 mimic was paired with BFP mRNA and the miR-122 sensor mRNA with iRFP mRNA (Figure 4A). Through StoichRNA, we sampled combinations of marker defined microRNA mimic and identified optimal performing conditions. We evaluated the microRNA sensor RNA with Ψ substitution or with unmodified uridine in separate StoichRNA experiments, allowing the operating conditions of the format to be assessed independently.

**Figure 4.**
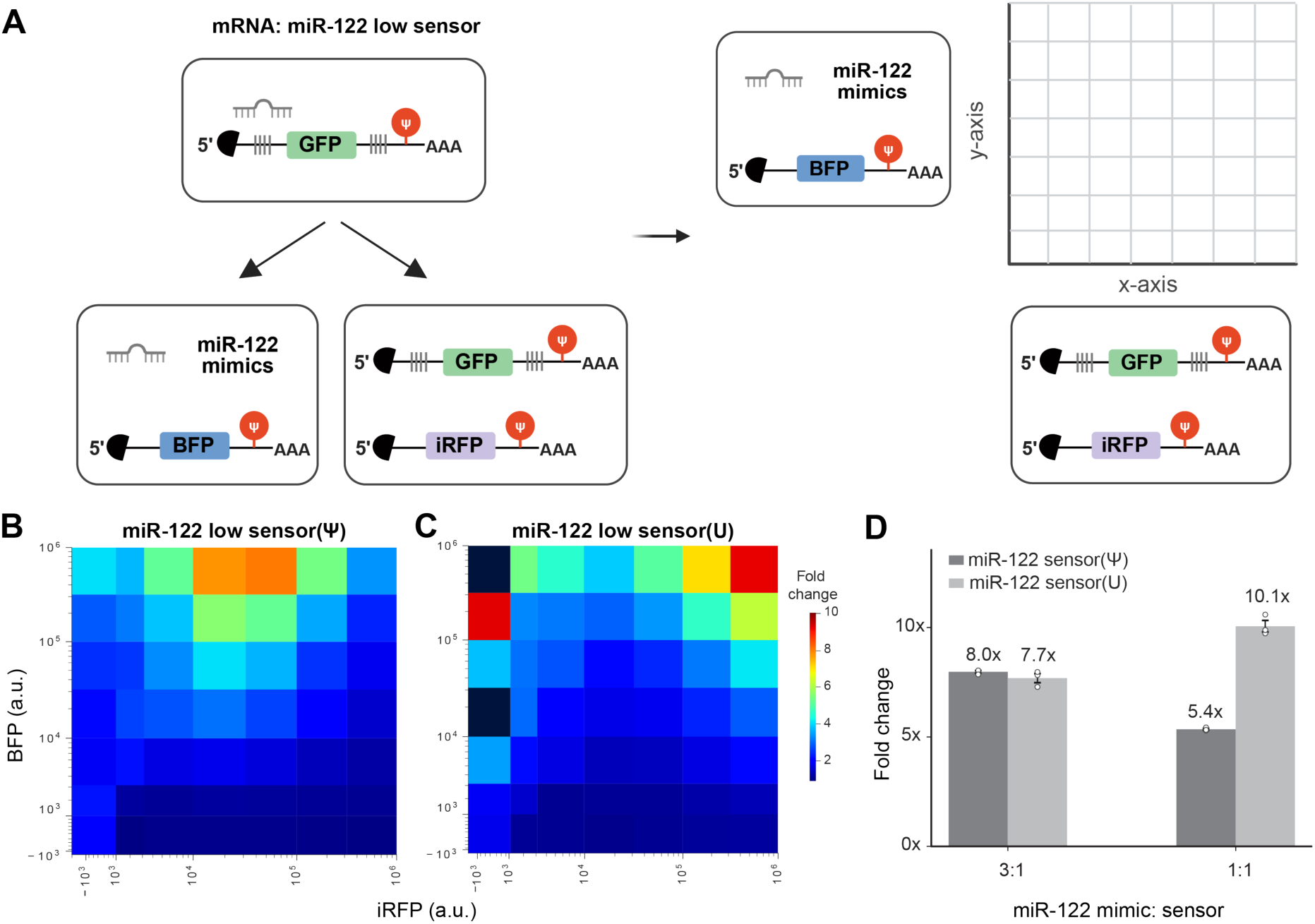
StoichRNA optimization of has-miR-122 low sensor. (A) Schematic of the hsa-miR-122 low sensor circuit: Ψ-substituted miR-122 low sensor with miR-122 mimics. Synthetic miR-122 mimics are co-delivered to titrate input levels and optimize sensor performance. In the StoichRNA experiment, miR-122 mimic delivery is mapped by BFP mRNA(Ψ) and sensor delivery is mapped by iRFP mRNA(Ψ). (B–C) Bin-analysis heatmaps of the miR-122 low sensor with Ψ (B) or U (C) substitution. (D) Histogram of the maximum fold change achieved at the optimal BFP: iRFP ratio for the Ψ sensor (3:1) and the U sensor (1:1). Ratios are based on BFP / iRFP mapped miR-122 mimic signal relative to mapped sensor signal.

We first binned cells by the two transfection-marker expression levels. The Ψ substituted miR-122-low sensor reached an 8x fold change in the highest miR mimic marker bin with intermediate sensor marker bin, while the unmodified sensor reached a comparable 10x fold change, clustered instead at high mimic-and sensor-marker fluorescence (Fig. 4B and C). To identify favorable relative input levels, we analyzed the data through trajectory sampling as described in figure 2. We found that the peak fold change of the miR-122-low sensor occurred at different marker signal ratios, with the Ψ-substituted sensor peaking around 3:1 miR-122 mimic: sensor ratio and the unmodified sensor around 1:1 (Figure 4D). Together, these results show that StoichRNA can identify optimal-preforming input to sensor ratio for RNA based sensing, with microRNA mimic dose as a tunable system input.

### StoichRNA enables optimization of a multi-component microRNA-high sensor across cell types

We next asked whether StoichRNA could be extended from the two-component miRNA-low sensor to a more complex multi-component RNA circuit. We used a miR-302-high sensor that converts high miR-302 activity into increased reporter expression (Figure 5A). In this architecture, the endoribonuclease CasE serves as an intermediate converter: miR-302 represses the CasE-encoding sensor RNA, thereby relieving CasE-mediated repression of the downstream reporter. The miR-302-high sensor therefore depends jointly on miR-302 activity and the relative levels of the sensor and reporter RNAs, creating a three-dimensional optimization problem well suited to StoichRNA.

**Figure 5.**
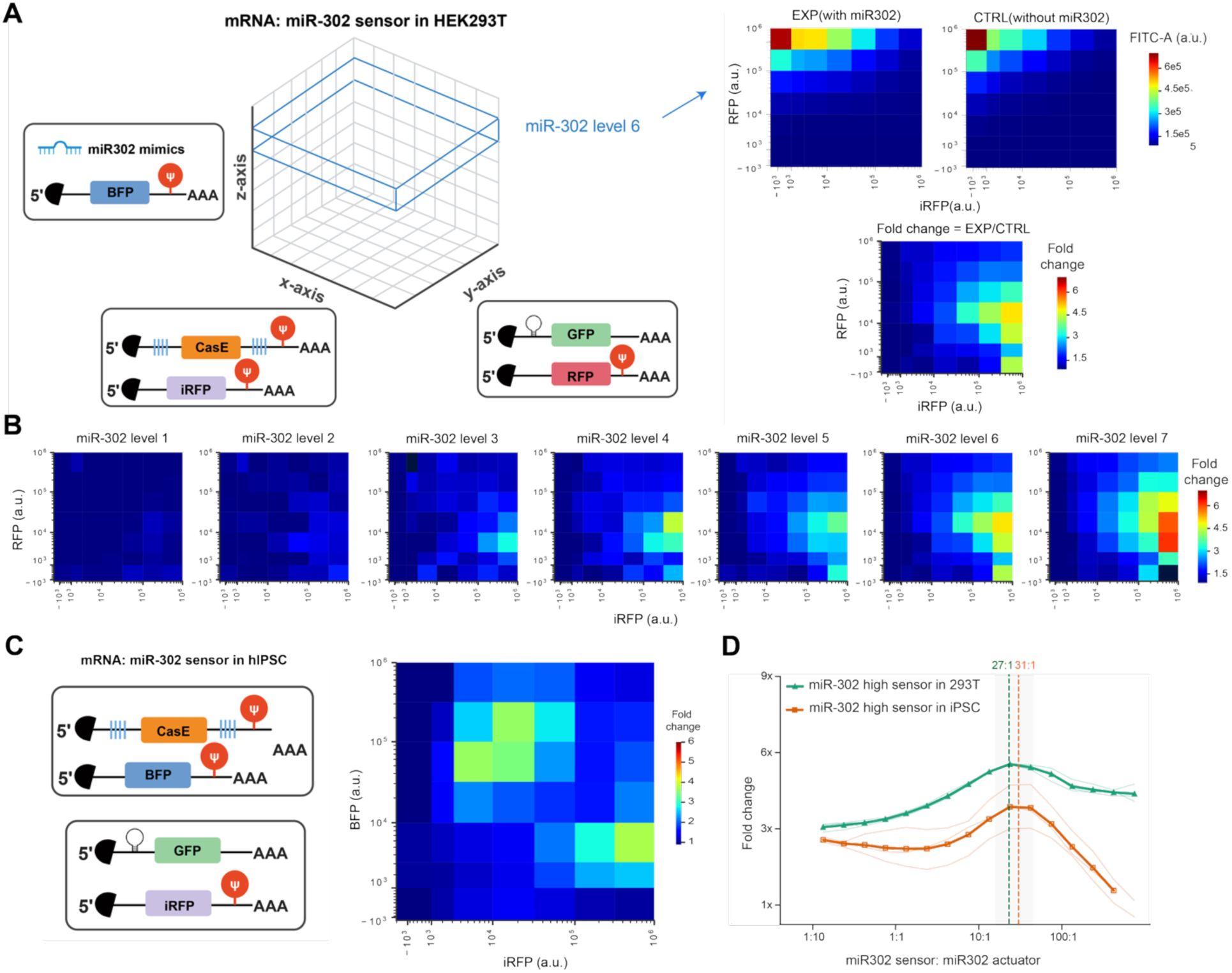
StoichRNA enables optimization of a multi-component microRNA-high sensor across cell types. (A) Visualization of four-dimensional miR-302 high-sensor StoichRNA data. The three modules of the sensor are each mapped by a separate fluorescent mRNA marker and organized into a binned 7 × 7 × 7 three-dimensional space; the mean GFP output is computed across all cells falling within each bin. The X and Y axes represent the sensor and reporter modules, respectively, and the Z axis represents miR-302 mimic level. At each miR-302 mimic level, the StoichRNA experiment comprises a paired EXP ( with miR-302 mimics) and CTRL ( without miR-302 mimics) sample, and the fold change heatmap is calculated as mean EXP fluorescence / mean CTRL fluorescence. Mimic level 6 is shown here as a representative example. (B) Fold change heatmaps of the Ψ-substituted miR-302 high sensor across all miR-302 mimic levels. (C) Schematic of the Ψ-substituted miR-302 high-sensor circuit in hiPSCs. The sensor module is paired with BFP(Ψ) and the miR-302 sensor reporter module is paired with iRFP(Ψ). (D) Fold change landscapes from trajectory sampling of the ratio space in HEK293T (green) and hiPSCs (orange). All the possible ratios through trajectories sampling are shown here. Across the sampled ratio space, the optimal miR-302 sensor: reporter ratio in HEK293T cells (27:1, green triangles; green dashed vertical line) and optimum identified in hiPSCs (31:1, orange squares; orange dashed vertical line).

Using the same general strategy used for the miR-122-low sensor, we used miR-302 mimics in HEK293T cells to span a gradient of miR-302 activities while simultaneously sampling sensor and actuator expression levels. For consistency with the stem-cell experiments below, the circuit used here was Ψ-substituted. StoichRNA generated a series of two-dimensional response landscapes across increasing miR-302 mimic levels, revealing a progressive increase in sensor activation as miR-302 input increased (Figure 5B). Within each miR-302 level, circuit performance varied substantially across the sensor–reporter expression space, demonstrating that miRNA input alone was insufficient to determine circuit output. Optimal-performing regions were enriched at intermediate-to-high sensor-marker levels together with lower relative actuator-marker levels.

We selected miR-302 because it is highly expressed in human induced pluripotent stem cells (hiPSCs)(27), allowing us to test whether the operating regimes identified in the HEK293T mimic model were preserved when the same circuit was delivered to an endogenous miRNA input in a distinct cellular context. To test this, we carried out StoichRNA profiling of the miR-302 high sensor directly in hiPSCs, which express miR-302 endogenously. Because no exogenous mimic was required, the circuit was profiled across the two-dimensional sensor–actuator space and compared with the corresponding high-miR-302 regions measured in HEK293T cells (Figure 5C). Both cell types were profiled using the same Ψ-substituted circuit, allowing the two datasets to be compared under matched RNA chemistry.

The hiPSC landscape exhibited a similar overall ratio-dependent response, with optimal-performing regions occurring within a comparable sensor–actuator regime. To compare the relative response profiles across cell types, we perform trajectory sampling with both HEK293T and hiPSC and then plotted the fold change landscape identified through a series of trajectory sampling, different expression levels of miR-302 mimics in HEK293T cell are pooled together so both of the system can be evaluated by sampling a 2D space. (Figure 5D). The HEK293T and hiPSC profiles showed similar overall trends with a clear peak performance overlapping, with max performance identified around 27:1 miR302 sensor: actuator in HEK293T (95% CI 22:1–33:1) and 31:1 in iPSC (24:1–40:1), The two optima sit within a ±1.7-fold margin, and the HEK293T optimum was able to retain 99% of hiPSC peak activation, indicating that the circuit’s stoichiometric behavior is preserved across cell types while circuit’s own fold change performance is cell-context-dependent. Together, these results show that StoichRNA can be used not only to optimize multi-component RNA circuits but also to assess how well operating regimes identified in one cellular context transfer to another.

### StoichRNA optimization of two-input RNA logic circuits

We next asked whether StoichRNA could be used to optimize multi-input RNA computation, where circuit performance depends simultaneously on several regulatory components. We constructed two logic circuits using the orthogonal endoribonucleases CasE and Csy4 as regulatory inputs (Figure 6A, F). First, we implemented a NOR gate in which CasE and Csy4 both act as repressors of a shared GFP output transcript carrying the corresponding CasE and Csy4 cognate sites (Figure 6A). Because our earlier results showed that Ψ substitution strongly reduced CasE-mediated repression, the CasE-responsive reporter was in vitro transcribed without Ψ substitution. Second, we implemented a CasE/Csy4 NAND gate comprising four mRNAs: one for CasE, one for Csy4, and two reporter mRNAs, each encoding GFP and carrying a cognate site for one of the two endoribonucleases (Figure 6F). Because either reporter can contribute to the output, fluorescence remains high when only one repressor is present and is reduced when both CasE and Csy4 are present, yielding a NOR gate.

**Figure 6.**
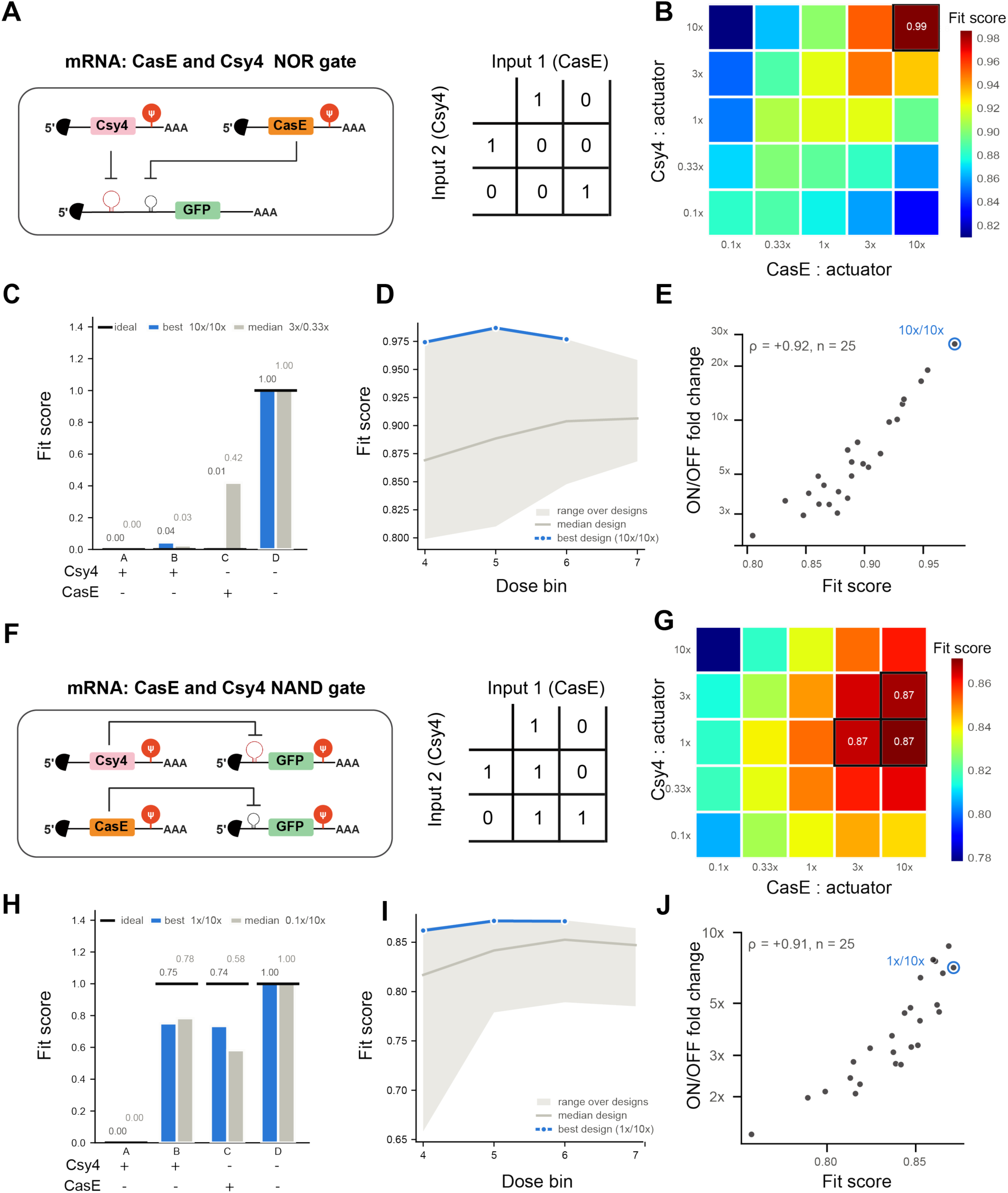
StoichRNA optimizations of two-input RNA NOR and NAND logic circuits. (A): Circuit shcematic and truth table of Endoribonuclease CasE and Csy4 RNA NOR gate (B, G): Gate fidelity heatmap across the stoichiometry scan space for NOR(B) and NAND(G) gate. Each cell is one design of Csy4: actuator (rows) × CasE: actuator (columns), each at 0.1×, 0.33×, 1×, 3× and 10×. Each cell’s fit score against the textbook truth table as fit score *=* 1 – mean |ŷ − t|, where ŷ is the log₁₀ normalized output level (0 = both inputs are true, 1 = neither inputs are true) and *t* is the target vector, (0, 0, 0, 1) for NOR and (0, 1, 1, 1) for NAND, where fit score close to 1 indicates the gate at selected ratio is close to a perfect textbook gate. Color bar is scaled within each gate’s fit score. Boxed cells are the best-scoring design based on fit score calculated; all ties are boxed. Reporter for both gate’s bin 5 is shown here as an example. (C, H): NOR(C) or NAND(H) gate measured truth tables for the best and median design, in the same log₁₀ normalized units. Black bars mark the ideal vector for each state, (0, 0, 0, 1) for NOR and (0, 1, 1, 1) for NAND. Values above each bar give the best (Black font, column in blue color) and median (light grey font, column in grey color) design. (D, I): Fit score of NOR(D) or NAND(I) gate against total dose along the actuator axis, for dose bins 4–7. The shaded band spans all 25 designs sampled, the grey line is the median design and the blue line the best design identified, same color as in E-F. (E, J): Fit score against operating window, for the NOR (I) and NAND (J) gates. Each point is one of the 25 designs (5 Csy4: actuator × 5 CasE: actuator ratios). The x axis is the fit score of panels C and D; the y axis is the ON/OFF fold change, defined as the smallest fold change between the state that should be ON and each state that should be OFF. The ringed point is the top-scoring design, labelled by its Csy4/CasE ratio (Blue). (F): Circuit shcematic and truth table of Endoribonuclease CasE and Csy4 RNA NAND gate

We applied StoichRNA to sample CasE, Csy4, and reporter-associated expression levels and evaluated circuit behavior across a three-dimensional marker-defined design space. For each gate, we examined 25 CasE:Csy4:actuator ratio combinations spanning a broad range of relative regulator levels (Figure 6B, G). Each computation was evaluated under four input conditions, denoted as [Input1, Input2] = [00, 10, 01, 11], with CasE and Csy4 as the two regulatory inputs correspondingly. We defined a fit score quantifying how closely a given design reproduced the desired truth table. For each sampled ratio, the normalized fluorescence vector across the four-input conditions was compared with desired output pattern, with a score of 1 indicating a perfect match. Using the input ordering shown in Figure 6, the ideal output vectors were (0, 0, 0, 1) for NOR and (0, 1, 1, 1) for NAND.

Across the 25 sampled designs, the NOR gate reached a maximum fit score of 0.99, with the highest-scoring region occurring at high relative CasE and Csy4 levels compared to the reporter (Figure 6C). The selected NOR design closely reproduced the ideal output pattern and outperformed the median design across the four input conditions (Figure 6D). Similarly, the NAND landscape revealed a maximum fit score of approximately 0.87 within a distinct region of CasE:Csy4 design space (Figure 6H) and the selected NAND design more closely matched the ideal NAND response than the median design (Figure 6I).

We next asked whether the optimized ratios remained effective across different overall reporter RNA doses. For both NOR and NAND gates, the best StoichRNA-selected designs maintained high fit scores across the tested actuator dose bins. The best design fit score of the NOR gate changes from 0.975 to 0.97, likewise, the best design of the NAND gate changes from 0.862 to 0.871, whereas the median designs showed lower overall performance, NOR median design fit score changes from 0.861 to 0.903, likewise, the best design of the NAND gate changes from 0.817 to 0.853 (Figure 6D, I). This indicates that StoichRNA can identify not only optimal-performing component ratios but also designs whose logic behavior is comparatively robust, indicating that the optimal ratio identified is a property of the design rather than of one particular delivery level of reporter.

To further verify the fidelity of the fit-score system, we considered a potential limitation. Because the four input conditions of each gate are normalized to their own min/max, the score may fail to capture the true dynamic range of the logic gate, which is central to its relevance for computation in biological systems as gate dynamic range sets the margin available for signal propagation when gates are layered in biological systems (23–25). A design correctly rendered as a NOR/NAND gate but weakly separated in magnitude would score as highly as one with a large dynamic window. We therefore asked whether designs selected by the fit-score also achieve the largest worst-case fold change, defined as the minimum fold change between the state that should be ON and each state that should be OFF. Score and worst-case fold change were strongly rank correlated across the 25 designs for both gates (Spearman ρ = 0.92 for NOR and 0.91 for NAND; both p < 1 × 10⁻⁹) (Fig. 6E, J), and this correlation held within every reporter dose bin examined (Figure 6D, I). Optimizing on fit-score therefore also selects for operating window, rather than trading one against the other.

Together, these results show that StoichRNA can efficiently search multidimensional RNA design space to identify optimal-performing and dose-robust configurations for multi-input logic computation.

## DISCUSSION

### StoichRNA enables systematic RNA circuit optimization

Component stoichiometry is a major determinant of RNA circuit performance, yet conventional co-transfection limits in testing one ratio per condition and makes systematic exploration costly. By deliberately manipulating the stochastic uptake of distinct transfection complexes, StoichRNA samples a wide range of stoichiometries within a single transfection, and the resulting data support multiple downstream analyses; binned heatmaps and trajectory sampling allows for systematic optimizations of a given system.

StoichRNA also captures how circuit behavior changes with nucleic acid format. In the CasE system, the DNA and RNA implementations differed substantially in both overall performance and in the optimal ratio between components, despite encoding the same circuit. Plausible contributors include differences in expression kinetics between transcribed and directly delivered RNA, and in the stability of each species. The practical implication is that a ratio optimized in one format cannot be assumed to transfer to the other.

One major limitation in the field is the uncertain transferability of circuits characterized in HEK293T cells to primary cell contexts. For the miR-302-high sensor, the optimal ratio identified by StoichRNA in HEK293T cells correlated strongly with that identified in hiPSCs, suggesting that ratios determined in a workhorse cell line may retain translational value in primary target cell types. This rests on a single sensor in one cell pair, however, and whether it holds for circuits with more components or stronger dependence on host RNA-binding factors remains to be tested.

### Differential effects of Ψ substitution across RNA regulatory mechanisms

Our results indicate that the effect of pseudouridine (Ψ) substitution is highly dependent on the regulatory mechanism. In the CasE OFF circuit, Ψ substitution of the target-containing reporter RNA strongly impaired repression, whereas replacing the reporter with unmodified RNA restored robust activity. This suggests that Ψ substitution can interfere with efficient CasE-mediated regulation. In contrast, the Csy4 OFF circuit remained functional when both regulator and reporter RNAs were Ψ-substituted, indicating that Csy4 activity is comparatively tolerant of this modification under the conditions tested. However, Figure 3 does not directly compare modified and unmodified Csy4 reporters, so a smaller effect cannot be excluded. For the miR-122 sensor, Ψ-substituted and unmodified RNAs achieved similar maximal repression, but their optimal response occurred at different marker-defined input-to-sensor ratios. Thus, Ψ substitution did not substantially reduce the achievable dynamic range of miRNA regulation but shifted the circuit operating regime. Together, these results show that Ψ substitution can have distinct consequences for different RNA regulators, ranging from strong functional impairment to preserved activity with altered optimal conditions, highlighting the need to evaluate RNA modification in a circuit-specific manner and to characterize and optimize RNA circuits in RNA format.

## ACKNOWLEDGEMENTS

We would like to acknowledge Edward Fusciello and Tengyuan Zhang for their help in circuit molecular cloning and IVT reactions. We would like to thank Lina Benjelloun for the discussions around the manuscript. We acknowledge the Institute of Chemical Imaging of Living Systems at Northeastern university for their resources around flow cytometry. We thank all Wang lab members for their feedback.

